# Computational modeling reveals cell-cycle dependent kinetics of H4K20 methylation states during Xenopus embryogenesis

**DOI:** 10.1101/2020.05.28.110684

**Authors:** Lea Schuh, Carolin Loos, Daniil Pokrovsky, Axel Imhof, Ralph Rupp, Carsten Marr

## Abstract

Histone modifications regulate chromatin architecture and thereby control gene expression. Rapid cell divisions and DNA replication however lead to a dilution of histone modifications and can thus affect chromatin mediated gene regulation So how does the cell-cycle shape the histone modification landscape, in particular during embryogenesis when a fast and precise control of cell-specific gene expression is required?

We addressed this question in vivo by manipulating the cell-cycle during early Xenopus laevis embryogenesis. The global distribution of un-, mono- di- and tri-methylated histone H4K20 was measured by mass spectrometry in normal and cell-cycle arrested embryos over time. Using multi-start maximum likelihood optimization and quantitative model selection, we found that three specific methylation rate constants were required to explain the measured H4K20 methylation state kinetics. Interestingly, demethylation was found to be redundant in the cycling embryos but essential in the cell-cycle arrested embryos.

Together, we present the first quantitative analysis of in vivo histone H4K20 methylation kinetics. Our computational model shows that demethylation is only essential for regulating H4K20 methylation kinetics in non-cycling cells. In rapidly dividing cells of early embryos, we predict that demethylation is dispensable, suggesting that cell-cycle mediated dilution of chromatin marks is an essential regulatory component for shaping the epigenetic landscape during early embryonic development.

## INTRODUCTION

All cells in our body contain the same genetic information encoded in the DNA. However, we are constituted out of many different cell types all performing their own specialized functions. Chromatin, mainly composed of DNA and histone octamers (two copies of histone H2A, H2B, H3 and H4 each), is an instructive DNA scaffold that aids extracting cell-specific information for gene expression. Histone tails are subject to various post-translational modifications such as methylation, acetylation, phosphorylation and ubiquitination (Bannister and Kouzarides, 2011), which play a fundamental role in altering chromatin accessibility. Dynamic regulation of gene expression is central for executing cell internal programs (proliferation, differentiation, etc.) and reacting to cell external signals with an appropriate response. articularly during development, where cells continuously divide and differentiate, a fast and economical control of gene expression is required. Histone modifications are believed to regulate the progression throughout development (Jambhekar et al., 2020). In Xenopus laevis, a model organism for developmental biology, stage-specific histone modifications have been observed during the transit from pluripotent to differentiated states, a process called epigenome maturation (Schneider et al., 2011). However, cells divide rapidly during early development. With each cell-cycle newly formed, largely unmodified histones are incorporated into the DNA leading to an overall dilution of most histone modifications (Jasencakova et al., 2010). How is the histone modification landscape shaped by the cell-cycle in vivo?

Histone methylation is known to play important roles in many biological processes (Greer and Shi, 2012) and its deregulation is linked to cancer and aging (Fraga et al., 2005; Klutstein et al., 2016). The methylation of lysine 20 on histone H4 (H4K20) is one of the most frequent lysine methylation sites (Jørgensen et al., 2013). It is evolutionarily conserved from Schizosaccharomyces pombe to humans (Lachner et al., 2004), and is known to have a strong cell-cycle dependence. H4K20 occurs in four different states, un-, mono-, di- and tri-methylation. Each methylation state plays a different functional role ranging from DNA damage repair (Sanders et al., 2004), over transcriptional regulation (Barski et al., 2007), chromatin condensation (Oda et al., 2009; Sanders et al., 2004), mitotic progression (Sakaguchi and Steward, 2007), and cell-cycle control (Schotta et al., 2008) to silencing of repetitive DNA and transposons (Schotta et al., 2004). H4K20me is regulated by three methyltransferases: KMT5A (also known as R-Set7) for mono-methylation (Xiao, 2005) and SUV4-20H1 and SUV4-20H2 for both di- and tri-methylation (Schotta et al., 2004). Whether there is a specificity of SUV4-20H1/2 for di- or tri-methylation is still debated (Schotta et al., 2008). The level of mono-methyltransferase KMT5A is cell-cycle dependent and its degradation in G1 phase leads to a decline of H4K20me1 in late G1 (Abbas et al., 2010; Centore et al., 2010; Zee et al., 2012). H4K20me1 reaches its lowest level in S phase while increasing in G2 phase and peaking during mitosis. Both H4K20me2 and H4K20me3 levels have also been found to be cell-cycle dependent though in a less dramatic fashion (Pesavento et al., 2008). The cell-cycle dependent presence of H4K20 methyltransferases allows H4K20me2 and H4K20me3 to be reestablished only after mitosis in the next cell-cycle (Jørgensen et al., 2013). For demethylation, unspecific enzymes such as HF8 are known (Feng et al., 2010), but their functional importance has recently been questioned (Alabert et al., 2020; Jørgensen et al., 2013; Reverón-Gómez et al., 2018). It has even been suggested that the loss of histone marks may occur only by dilution during chromatin replication than by active removal (Jadhav et al., 2020).

To address the role of the cell-cycle for epigenome maturation in Xenopus development, we have measured histone modification proportions in sibling embryo populations, which either proliferate or are arrested at the G1/S transition. Using quantitative mass spectrometry data for H4K20 we compared over 200 model hypotheses describing H4K20me kinetics in the cycling and cell-cycle arrested population. With only a few assumptions, our computational model is able to explain H4K20me kinetics, retrieves correct cell-cycle durations and known cell-cycle dependencies of H4K20me. Furthermore, our approach allows us to estimate cell numbers over time and reveals the importance of three specific methylation rate constants and a shared demethylation rate constant which is essential to establish the observed histone modification profile in the cell-cycle arrested but redundant in the cycling population of Xenopus embryos.

## RESULTS

### Cell-cycle arrest changes H4K20me patterns during Xenopus embryogenesis

After in vitro fertilization of a Xenopus oocyte, cells rapidly divide in a state of transcriptional quiescence up to 5.5 hours post fertilization (hpf) (Heasman, 2006). Only then a regular zygotic cell-cycle containing G1 and G2 phases is initiated (Newport and Kirschner, 1982). To identify how H4K20 methylation (H4K20me) is shaped by cell-cycle, we compared a population of normal Xenopus embryos (from now on called ‘mock’) with a cell-cycle arrested population. For this, half of the embryos were continuously incubated with hydroxyurea/aphidicolin (from now on called ‘HUA’) from gastrulation onwards (11 hpf). This treatment arrests cells at the G1/S boundary and is compatible with embryonic development (Harris and Hartenstein, 1991). Mass spectrometry measurements of H4K20me states, averaging over all cells in the embryos and all histones in the cells, were conducted at 14.75, 19.75, 27.5 and 40 hpf corresponding to late gastrula (NF13), neurula (NF18), tailbud (NF25) and tadpole (NF32) stages, respectively (Figure 1A). H4K20me proportions of mock and HUA showed clear differences across three biological replicates in all four H4K20me states (Figure 1B). In HUA, methylation accumulates in the di- and tri-methylation states in comparison to mock. There, newly synthesized and unmodified histones are incorporated upon DNA replication, leading to an overall dilution of H4K20me and hence a higher proportion of lowly methylated un- and mono-methylated states.

**Figure 1.**
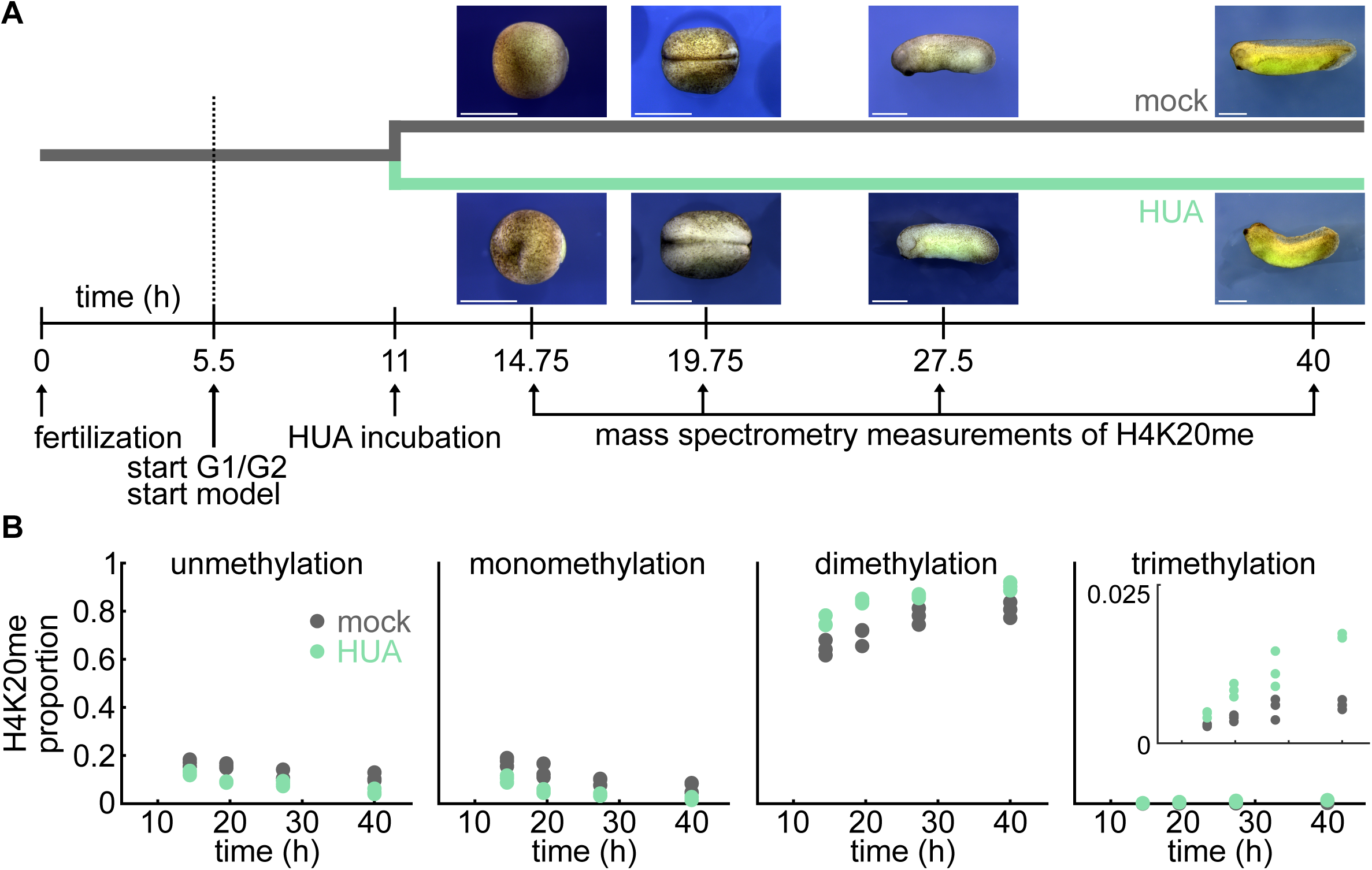
H4K20 methylation kinetics during Xenopus embryogenesis are altered upon HUA induced cell-cycle arrest. **(A)** Xenopus eggs are fertilized in vitro at time point 0. For the next five hours post fertilization (hpf), the embryonic cell-cycle consists of S and M phases only. At 5.5 hpf, G1 and G2 phases appear. At 11 hpf, half of the embryos are incubated with hydroxyurea/aphidicolin (HUA), arresting cells at the G1/S boundary. Mass spectrometry measurements of H4K20 methylation (H4K20me) are performed at 14.75, 19.75, 27.5 and 40 hpf in embryos with dividing (mock) or non-dividing cells (HUA). HUA incubated embryos are viable and visually remarkably similar to mock embryos (scale bar 1mm). **(B)** H4K20me kinetics differ between mock (gray) and HUA treated (green) embryo populations. In HUA H4K20 un- and mono-methylation is decreased while H4K20 di- and tri-methylation (see inset) is increased.

### Specific methylation rate constants are necessary to explain H4K20me in mock embryogenesis while demethylation is not essential

To identify how H4K20me kinetics are shaped by cell-cycle, we defined models for untreated embryos (mock) and fitted them to the data (see Methods). Mock models are composed of four H4K20me states corresponding to un- (me0), mono- (me1), di- (me2) and tri-methylated (me3) H4K20, allowing for successive methylation and demethylation with rates m_i_ and d_i_, i ∈ {1,2,3} respectively (see Figure 2A and Methods for a detailed model description). For mock, where the cells undergo cell division with a cell-cycle duration c, unmethylated histones are incorporated into replicating DNA leading to a continuous dilution of H4K20me. Considering methylation proportions (defined as the frequency of a particular methylation state divided by the sum of all methylation states as measured by mass spectrometry), cell-cycle results in an overall increase of unmethylated H4K20 mediated by an outflow of H4K20me states with rate g(t) = ln(2)/c(t), where c(t) is the cell-cycle duration c as a function of experiment time t (see Figure 2A and Methods). The most general model is parameterized with six rate constants, where a rate constant is defined as the proportion of H4K20 in a particular methylation state being methylated/demethylated per hour (h^-1^), and contains three rate constants for methylation m_1_, m_2_, m_3_ and three rate constants for demethylation d_1_, d_2_, d_3_ (Figure 2B, rightmost model). However, we also considered models with less parameters: rate constants shared between two or more reactions are termed ‘shared methylation/demethylation rate constants’ (Figure 2B, gray) and rate constants specific to one reaction are termed ‘specific methylation/demethylation rate constants’ (Figure 2B, colored). Intrigued by the question whether demethylation is important for methylation kinetics at all (as its existence was recently challenged at least for histone H3 lysine 27 tri-methylation (Reverón-Gómez et al., 2018), we also considered 5 models without demethylation. In total, the 30 models we consider comprise between 1 and 6 rate constants (Figure 2B and Methods). In addition to the rate constants, we inferred another 4 model parameters: 3 initial H4K20me proportions at 5.5 hpf (denoted as me0_0_, me1_0_, me2_0_, me3_0_ with me0_0_=1-me1_0_-me2_0_-me3_0_), and one noise parameter σ, determining the width of the Laplacian noise distribution (Methods). As we were interested in H4K20me kinetics under the influence of the cell-cycle, we started our mock model at 5.5 hpf (Figure 1A), when a regular zygotic cell-cycle with G1/G2 phases is initiated (Newport and Kirschner, 1982). Since cell cycle has been shown to vary substantially with embryonic age, we considered 6 different cell-cycle functions c(t) to model cell cycle over the experiment time t: constant, linearly increasing, or gradually plateauing (using a scaled Hill function with Hill coefficient 1 and offset) (Figure 2C). The number of model parameters for the cell-cycle functions varied from 1 (for the constant cell-cycle function) to 3 parameters (for the gradually plateauing cell-cycle function) (Methods). We performed multi-start maximum likelihood optimization and model selection on 180 models (30 models times 6 different cell-cycle functions). Including prior biological knowledge about the short cell-cycle at 5.5 hpf of ∼30 min (Anderson et al., 2017; Gelens et al., 2015), we found that only one of the six tested cell-cycle functions was able to predict a biologically meaningful average cell-cycle duration of around 8 hours: a constrained scaled Hill function with Hill coefficient 1 and offset 0.5, c(t) = 0.5 + b(t/(b+t)) (Supplementary Information). All models with other cell-cycle functions estimated average cell-cycle durations of at least 70 hours. Using a constrained scaled Hill function, we found 12 models that outperformed other models with a BIC (Bayesian Information Criterion) difference of ΔBIC>10, which is considered to be an appropriate threshold for model rejection (Kass and Raftery, 1995) (Figure 2D). The two best models (with ΔBIC=0) show specificity in tri-methylation and shared rate constants for mono- and di-methylation. Overall, the best models with and without demethylation showed specificity in either all three methylation rate constants or only in the tri-methylation rate constant. Varying numbers of demethylation rate constants were possible, ranging from 0 to 3. Fits to these 12 top models were able to capture the kinetics underlying H4K20me during mock embryogenesis (Figure 2E). Together, we found that either three specific methylation rate constants or one specific tri-methylation rate constant were necessary to explain the data from untreated embryos and that active demethylation was not required.

**Figure 2.**
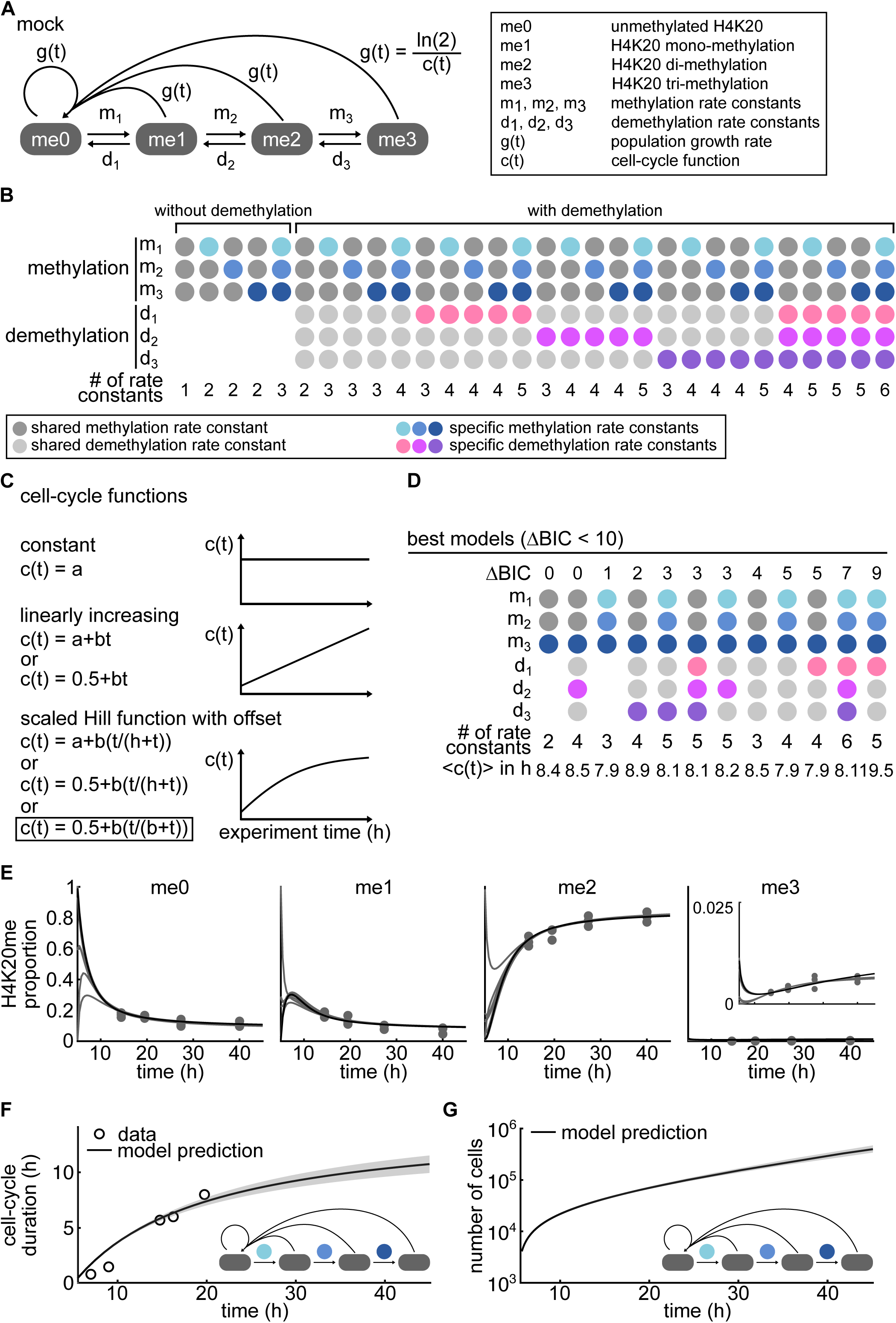
Demethylation is redundant to explain data of cycling mock cells. **(A)** Model of cycling mock population composed of four H4K20 states: un- (me0), mono- (me1), di- (me2) and tri-methylation (me3). m_1_, m_2_, m_3_ represent the mono-, di- and tri-methylation rate constants, and d_1_, d_2_, d_3_ represent the demethylation rate constants. An overall dilution of methylation happens due to cell division, parametrized with population growth rate g(t), which is dependent on the cell-cycle function c(t). **(B)** All possible parameter combinations result in 5 models without demethylation and 25 models with demethylation. Rate constants specific to a particular methylation or demethylation step are indicated in color, rate constants shared between methylation or demethylation steps are shown in gray. The number of rate constants ranges between 1 for the simplest model with no demethylation and shared methylation rate constant and 6 for the most complex model, where each methylation and demethylation rate constant is specific. **(C)** Only a constrained scaled Hill function with Hill coefficient 1 and offset 0.5 gives a cell-cycle duration in the expected range of 8 hours (marked by the black box). All other cell-cycle functions c(t) predicted average cell-cycle durations of at least 70 hours, which is biologically not meaningful and reflects a population of non-cycling cells. **(D)** The 12 best performing models are ordered by increasing BIC (Bayesian Information Criterion). All models with ΔBIC < 10 require either three specific methylation rate constants (m_1_, m_2_, m_3_) or a specific tri-methylation rate constant. However, if present, demethylation may take on any of the 5 possible rate constant combinations. The best performing models without demethylation perform similarly well as the best performing models with demethylation (ΔBIC=0 and 1). The estimated average cell-cycle duration <c(t)> is in a biologically realistic range of around 8 hours. **(E)** All 12 best performing models fit the data. The model with three specific methylation rate constants but with no demethylation is shown in black. **(F)** Model prediction of the cell-cycle duration (median, 25th and 75th percentiles of MCMC chain of the cell-cycle parameter of the model with three specific methylation rate constants but with no demethylation (inset)) agrees with experimental measurements of different papers. **(G)** The model with three specific methylation rate constants but with no demethylation (inset) predicts an increase of cell numbers from roughly 20,000 cells after 10h to 300,000 cells after 40h (using the median, 25th and 75th percentiles of the MCMC samples of the cell-cycle parameter of the model with three specific methylation rate constants but with no demethylation (inset)) in a developing Xenopus embryo.

### Validation of mock model by comparing cell-cycle durations to experimental data

We validated one of the best performing models by comparing it to the average cell-cycle durations experimentally measured in Xenopus neural progenitors at various developmental stages (Graham and Morgan, 1966; Sabherwal et al., 2014; Thuret et al., 2015). The cell-cycle durations from the mock model with three specific methylation rate constants but no demethylation (Figure 2F, inset) showed good agreement with measured cell-cycle durations (Figure 2F). Using this model, we can also predict the absolute number of cells within a developing embryo, which is experimentally challenging. For the same model (Figure 2G, inset), the number of cells was predicted to rise exponentially from roughly 20,000 cells after 10 hours to 300,000 cells after 40 hours (Figure 2G and Methods).

### Specific methylation rate constants and demethylation are necessary to model H4K20me in HUA embryogenesis

In contrast to mock, methylated H4K20 is not diluted in the cell-cycle arrested HUA embryo population. We thus modeled HUA with the same set of reactions, however, without a cell-cycle function g(t)=0 (Figure 3A). Similarly to the mock model we performed multi-start maximum likelihood optimization and model selection on 30 HUA models with and without demethylation. We found that the five best performing models (with ΔBIC<10) all required three specific methylation rate constants and demethylation (Figure 3B). The number of demethylation rate constants varied between 0 and 3 (Figure 3B). The single best performing HUA model without demethylation (rightmost model in Figure 3B) was substantially outperformed by the HUA models with demethylation (ΔBIC=13) suggesting that demethylation was essential to explain the HUA data. The model fits of the five best HUA models were able to capture the kinetics underlying H4K20me during HUA embryogenesis (Figure 3C). Together, we found that three specific methylation rate constants were necessary to explain the HUA data and that demethylation was essential.

**Figure 3.**
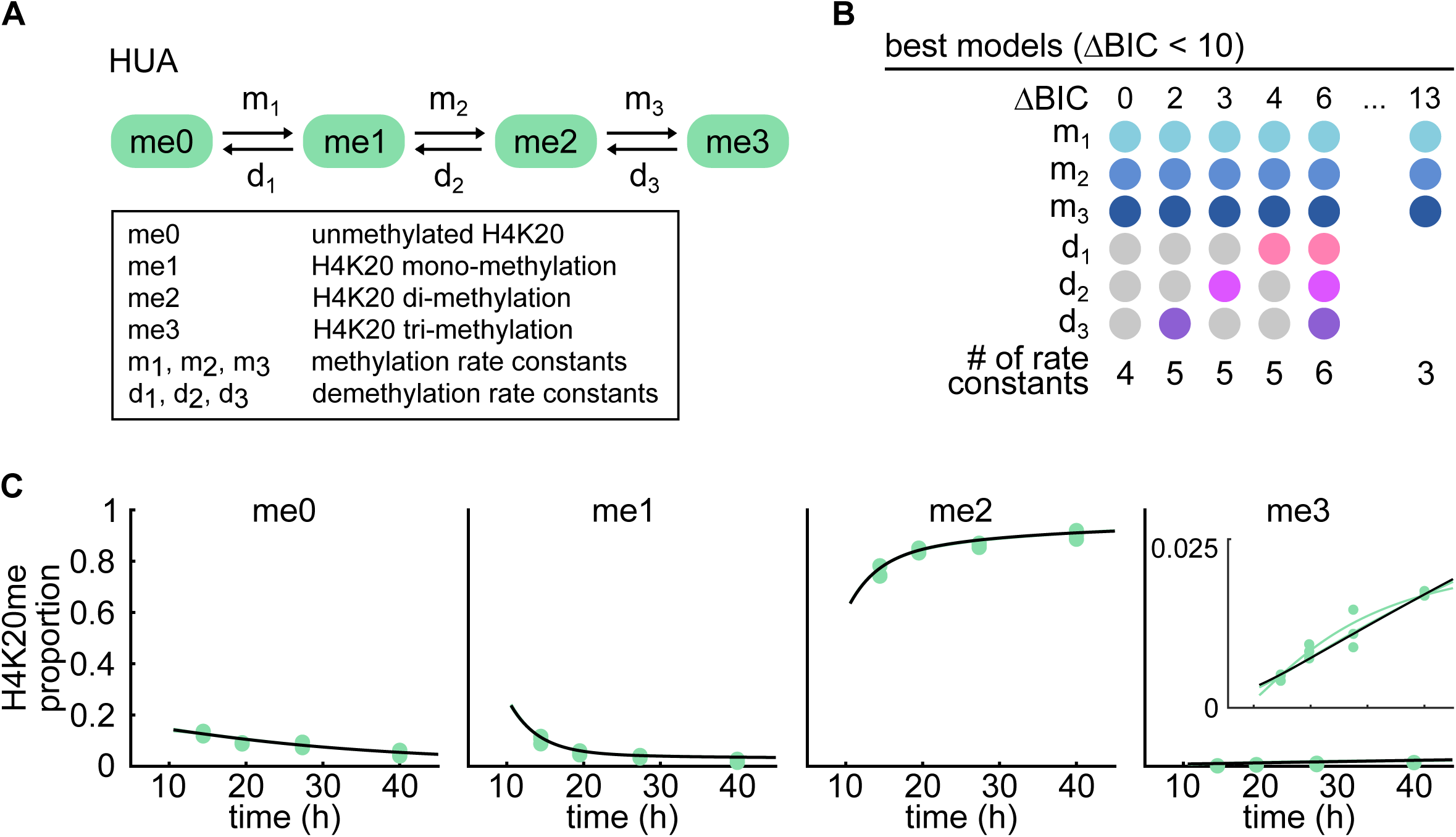
Demethylation is essential to explain data of cell-cycle arrested HUA cells. **(A)** Model of cell-cycle arrested HUA population. In contrast to the mock model (Figure 2A), the HUA cells do not divide (g(t) = 0) and no dilution of methylated H4K20 is required. **(B)** The 5 best performing HUA models with ΔBIC<10 all require 3 specific methylation rate constants (m_1_, m_2_, m_3_) and demethylation. However, demethylation may take on any of the 5 possible rate constant combinations. The single best performing HUA model without demethylation (right) is outperformed by the HUA models with demethylation (ΔBIC=13). **(C)** Model fits of top 5 HUA models with demethylation overlap strongly and show the ability to explain the HUA data. The best performing model is highlighted in black.

### Joint model is able to retrieve cell-cycle dependence of H4K20me and finds demethylation to be essential in HUA but redundant in mock

The models performing best in mock and HUA required three specific methylation rate constants and were indecisive about demethylation ranging from no demethylation over one shared to three specific demethylation rate constants (Figure 2D and Figure 3B). To determine which rates are substantially different between the two Xenopus populations we considered these findings and devised a joint model considering mock and HUA data simultaneously. For the most general hypothesis (Figure 4A), we allowed for three mock- and three HUA-specific methylation rate constants (visualized by a half gray and half green dot for m_1_, m_2_ or m_3_ in Figure 4B). We also allowed for joint methylation rate constants shared between specific mock and HUA methylation steps (visualized as an orange dot for m_1_, m_2_ or m_3_ in Figure 4B) reducing the number of parameters. As demethylation was not necessary to explain the mock data and one demethylation rate constant shared between methylation steps was sufficient for HUA, we here restricted demethylation to the simplest case of at most one shared demethylation rate constant d_mock_ and d_HUA_ (Figure 4A). We allowed for mock- and HUA-specific demethylation rate constants (visualized again by a half gray and half green dot for d in Figure 4B) or a joint demethylation rate constant for mock and HUA (visualized as an orange dot for d in Figure 4B). Furthermore, as a constrained scaled Hill function with Hill coefficient 1 and offset 0.5 was the only function that led to biolofgically meaningful cell-cycle durations (see above and Figure 2C and 2F) we did not consider different cell cycle functions thereby reducing the set of possible models to 8 joint models without demethylation (Figure 4B left), 16 joint models with demethylation and 2 × 8 models with demethylation in either mock or HUA (Figure 4B right). To identify joint models that are able to explain our measured data, we again fitted the models using multi-start maximum likelihood optimization and model selection.

**Figure 4.**
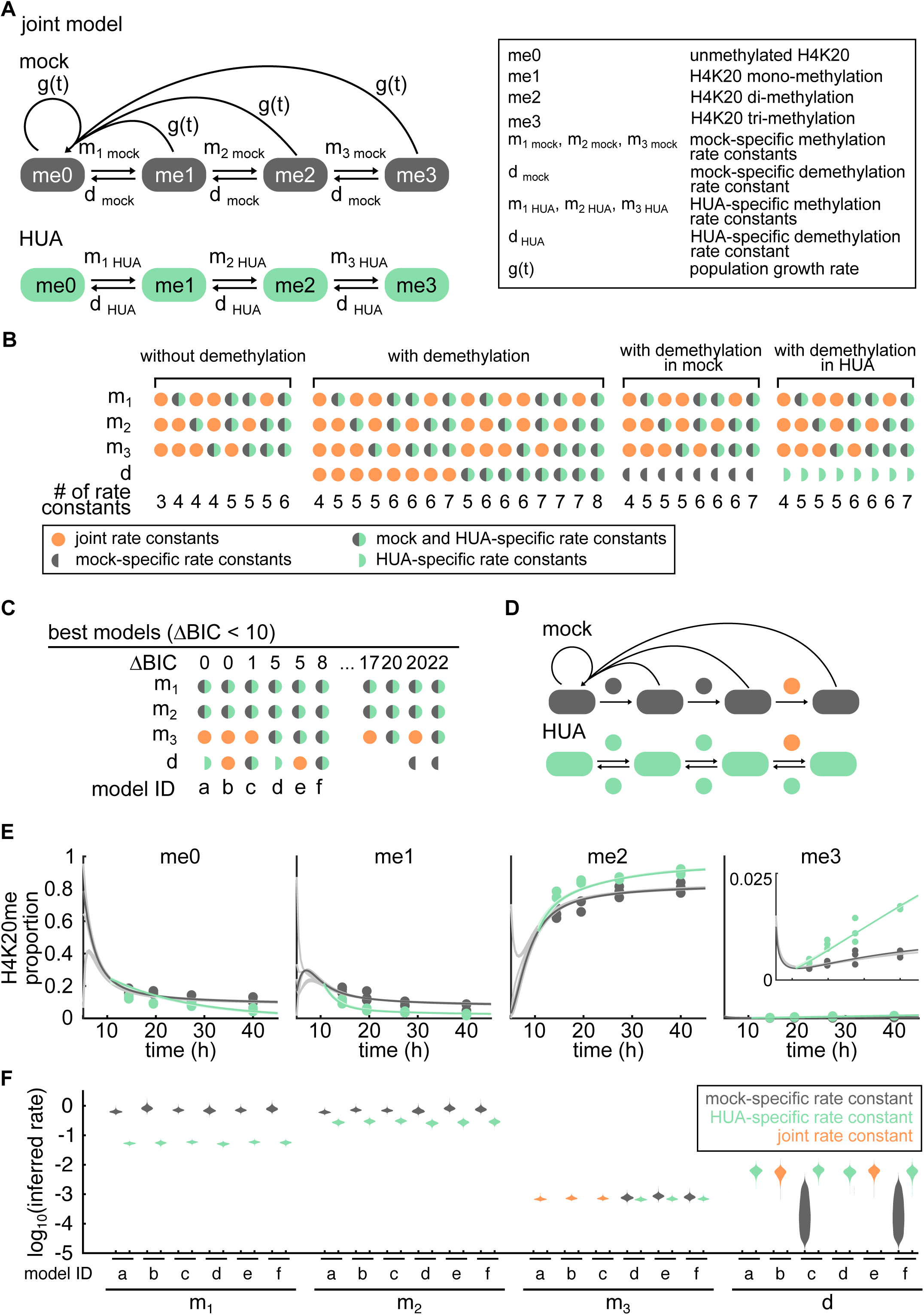
Joint computational modeling allows direct comparisons between mock and HUA rate constants and reveals that demethylation is overshadowed by HUA. **(A)** Joint model allows for three methylation and one demethylation rate constants for both mock and HUA as suggested by the best models for mock and HUA. **(B)** We fit 16 models with demethylation and 8 models each for without demethylation in mock and/or HUA to the joint data to infer mock- and HUA-specific rate constants. The joint rate constants of mock and HUA are shown in orange, the rate constants present in both the mock and HUA models but taking on mock- and HUA specific values are indicated in gray/green, the rate constants only present in the mock or HUA model are shown in gray and green half-circles, respectively. The model structure of the most complex of models is shown in (A). The number of rate constants ranges between 3 and 8. **(C)** The best performing models on the combined data set are ordered according to their BIC value. All models require HUA-specific mono- and di-methylation rate constants, but are indecisive about tri-methylation and demethylation. Joint models where demethylation is present in either only HUA or in both mock and HUA perform equally well. Joint models where demethylation is not present in either only HUA or in both mock and HUA perform considerably worse. Model IDs of all considerably best performing models are given (a-f). **(D)** Model structure of the simplest best performing joint model with demethylation in only HUA (model a). **(E)** All best performing joint models are able to explain both the mock and HUA data. The estimated initial conditions vary between the models. Joint model a is highlighted. **(F)** The violin plots of the marginal distributions of all best performing joint models show high consistency between the estimated methylation and demethylation rate constants. HUA-specific mono- and di-methylation rate constants are considerably decreased. Tri-methylation rate constants between mock and HUA have strongly overlapping marginal distributions. Demethylation seems to be dominated by the HUA population and is negligible in the mock population if a mock-specific demethylation rate is allowed.

All 6 best performing joint models (ΔBIC<10) required mock- and HUA-specific mono- and di-methylation rate constants (Figure 4C). However, they were not conclusive about tri-methylation and, if present, demethylation rate constants (Figure 4C). Specificity in one or more rate constants highlights that the differences in H4K20me proportions of mock and HUA are not explicable by the missing cell-cycle alone but that the overall H4K20me kinetics are cell-cycle dependent. The model structure of the best performing joint model (model a) is shown in Figure 4D. Interestingly, joint models with demethylation in HUA only (models a and d in Figure 4C) performed just as well as joint models with demethylation in both HUA and mock (models b, c, e and f in Figure 4C) while joint models without demethylation (ΔBIC=17 and 20) and joint models with demethylation in mock only (ΔBIC = 20 and 22) were substantially outperformed. This suggested that demethylation was essential for HUA only, in accordance with the results from the separate models (Figure 2D and Figure 3B).

The top 6 joint models (models a-f in Figure 4C) showed good overall agreement with mock and HUA data (Figure 4E) and strongly consistent rate constants (Figure 4F). We determined the marginal distributions for all rate constants by MCMC sampling, where the credibility ranges are the 25^th^ and 75^th^ percentiles of the marginal distributions (see Methods).P articularly interesting were the strong discrepancies between mono- and di-methylation rate constants for mock and HUA, decreasing 10-fold and 2-fold, respectively (Figure 4F). The mock-specific mono- and di-methylation rate constants of the top 6 joint models had overlapping credibility ranges suggesting that for mock, a shared rate constant for mono- and di-methylation would suffice (Figure 4F). Similarly, mock- and HUA-specific tri-methylation rate constants show overlapping credibility ranges suggesting that a joint tri-methylation rate constant would suffice. In joint models with demethylation (models c and f) we found the mock-specific demethylation rate constants to take on very small values while the HUA-specific demethylation rate constants were small but substantially larger: for model f the median mock-specific demethylation rate constant was estimated to be 2.0 10^−4^ (with 0.8-5.8 10^−4^ credibility range), while the HUA-specific demethylation rate constant was estimated to be 5.8 10^−3^ (4.7-6.9 10^−3^) (Figure 4F).

## DISCUSSION

We developed a computational model to investigate how H4K20me states are shaped by cell-cycle during Xenopus embryogenesis. Our findings support the notion that cell-cycle influences methylation kinetics and suggests that demethylation is redundant in cycling but essential in non-cycling cells.

### Active demethylation is redundant for mock but essential for HUA treated embryos

Comparing mock and HUA models with and without demethylation we found that demethylation is redundant in mock but essential undercell cycle arrest (Figure 2D and Figure 3B). We verified these findings with a joint model where the mock and HUA data is modeled simultaneously. Interestingly, the joint demethylation rate constants (in models b and e) were estimated to similar values as HUA-specific demethylation rate constants (for model b the joint demethylation rate constant was estimated to be 5.9 10^−3^ (4.8-7.0 10^−3^), for model f the the HUA-specific demethylation rate constant was estimated to be 5.8 10^−3^ (4.7-6.9 10^−3^)). This suggests that joint demethylation rate constants are completely overshadowed by the HUA model, strengthening the hypothesis that demethylation is redundant in mock but essential in HUA. Biologically, this could mean that while demethylases are present during embryogenesis, their effect in cycling cells is minute due to an overall dilution by unmodified histones. Only when cells stop to cycle (as modelled with the HUA treatment in our approach) demethylation kicks in and stabilizes posttranslational modifications specifically, thereby potentially driving differentiation.

### HUA-specific mono- and di-methylation are strongly decreased with respect to mock

Our findings can be interpreted in light of the current knowledge on methyltransferases. The mono-methyltransferase KMT5A (PR-Set7) was found to be cell-cycle dependent, getting degraded by the proteasome in G1 phase (Abbas et al., 2010; Centore et al., 2010). In the absence of KMT5A, mono-methylation might be compensated by SUV4-20H1/2 but with lower activity (Southall et al., 2013; Yang et al., 2008). HUA treatment blocks the cell-cycle at the G1/S boundary, suggesting that none to little KMT5A is present in HUA to mono-methylate H4K20. This is reflected by a 10-fold decrease in the HUA-specific mono-methylation rate constant in all best performing joint models (Figure 4F). As HUA-specific mono-methylation rate constants were necessary to explain the data (see Figure 4C), the joint model is able to retrieve this known cell-cycle dependence of H4K20me. H4K20me2 is also regulated in a cell-cycle dependent manner, however peaking in G1 phase (Pesavento et al., 2008). In contrast, all best performing joint models estimate the HUA-specific di-methylation rate constants to be decreased 2-fold in comparison to the mock-specific di-methylation (Figure 4F). We hypothesize this unexpected decrease of HUA-specific di-methylation to be due to either compensatory effects of SUV4-20H1/2, when the enzymes additionally mono-methylate H4K20, or so far unknown effects.

### Computational approach suggests shared rate constant for mono- and di-methylation in mock

The separate model for mock identified either only tri-methylation or all three methylation steps to be specific (Figure 2D). The joint model reflects the same specificities regarding methylation in mock. Even though the joint model allows for specificity in all three methylation steps, the credibility ranges of mock-specific mono- and di-methylation rate constants in the joint models overlap (Figure 4F left). This suggests that a shared rate constant for mock mono- and di-methylation would suffice resulting in a mock model with specificity in tri-methylation only. However, in the joint models the HUA-specific mono- and di-methylation rate constants have non-overlapping credible ranges with respect to the mock-specific rate constants (10-fold and 2-fold decrease) nor to each other. Under the assumption that the mock and HUA models are based on the same model structure, allowing for three specific methylation rate constants in the joint model was thus necessary (Figure 4F) to resolve these differences. We cannot infer from specificities in rate constants to specificities in the enzymatic activity of methyltransferases. However, current research hypothesizes that there exist three different methyltransferases: KMT5A performing mono-methylation (Xiao, 2005) and SUV4-20H1/2 performing both di- and tri-methylation (Schotta et al., 2004). Whether there exists a specificity of SUV4-20H1/2 for di- or tri-methylation is still debated (Schotta et al., 2008).

### Joint model suggests that tri-methylation is not necessarily cell-cycle dependent

All three joint models with specificity in tri-methylation (models d, e and f) result in slightly lower BIC values (Figure 4C), which is likely due to an increased penalization term for an additional estimated parameter and not due to a decreased likelihood. The estimated tri-methylation rate constants are small (on the order of 10^−3^) and the credible ranges for mock- and HUA-specific tri-methylation overlap in all three joint models suggesting that a joint tri-methylation rate constant would suffice. When we interpret differences in HUA and mock rates as indications for cell cycle dependent rates, we find no evidence for cell-cycle dependence for H4K20 tri-methylation. To clarify if the corresponding enzymes are indeed homogeneously expressed is up to further research.

Together, we provided a computational framework to analyze how H4K20me states are shaped by the cell-cycle throughout early development. We model the evolution of H4K20me states during embryogenesis in two Xenopus populations - one population with a regular cell-cycle, the other population with an arrested cell-cycle. We found that our model is able to retrieve the known cell-cycle dependence of H4K20me, where mono- and di-methylation rate constants are substantially decreased in HUA. Additionally, our models - both the mock and HUA models as well as the joint model - propose that demethylation is only essential in the cell-cycle arrested HUA population while redundant in the cycling cells of the mock population.

## ACKNOWLEDGEMENTS

L.S. was funded by the BMBF project TIDY (031 0170B). L.S. is especially grateful to the Technical University of Munich’s Department of Mathematics, whose generous Entrepreneurial Award (within the rogram “Global Challenges for Women in Math Science”) contributed to the completion of this project. D.P., A.I. and R.R. were funded by the Deutsche Forschungsgemeinschaft (DFG, German Research Foundation) - roject ID213249687 - SFB 1064. Animal work has been conducted in accordance with Deutsches Tierschutzgesetz; Xenopus experiments were approved by the Government of Oberbayern.

## AUTHOR CONTRIBUTIONS

Conceptualization, A.I., R.R. and C.M.; Methodology,L.S., C.L.; Software,L.S., C.L. Validation, C.L.; Formal Analysis,L.S.; Investigation,L.S., D.P.; Resources, A.I., R.R., C.M.; Data Curation, D.P.; Writing - Original Draft,L.S.; Visualization,L.S.; Supervision, A.I., R.R., C.L. and C.M.; Project Administration, A.I., R.R. and C.M.; Funding Acquisition, A.I., R.R. and C.M.

## MATERIALS AND METHODS

### Embryos handling and HUA treatment

Xenopus laevis eggs were collected, in vitro fertilized and handled by standard methods (Sive et al., 2000). The staging was done according to Nieukoop and Faber (Nieuwkoop and Faber, 1994). When embryos reached the desired stage (NF10.5), they were separated into two groups and incubated continuously into either HUA or mock solutions in parallel. HUA solution: 20mM Hydroxyurea (USBiological, H9120) and 150µM Aphidicolin (BioViotica, BVT-0307) in 0.1x MBS solution (Harris and Hartenstein, 1991). Mock solution: 2% DMSO (dissolvent for Aphidicolin) in 0.1x MBS solution. The embryos were collected at the four developmental stages (NF13, NF18, NF25 and NF32) for the mass spectrometry analysis.

### Nuclear histone extraction

Around 50 to 200 embryos developed to desired stages (NF13, NF18, NF25 NF32). They were harvested and histone proteins were purified by acid extraction from nuclei (Schneider et al., 2011, okrovsky et al., submitted). Each developmental stage is represented by three biological replicates. Each biological replicate derived from a different mating pair.

### Mass spectrometry sample preparation

The pellet from the nuclear histone extraction was dissolved in an appropriate amount of Lämmli Buffer to reach 1.37 10^6^ nuclei/µl in each sample. 15µ were loaded on an 8-16% gradient SDS-PAGE gel (SERVA ot V140115-1) and stained with Coomassie Blue to visualize the histone bands. Histone bands were excised and propionylated (as described in (Villar-Garea et al., 2012)). As an internal and inter-sample control, a library consisting of heavy-labelled peptides mimicking H4K20 methylation states which contain a heavy Arginine (R10 peptides) was used (product of JPT company). R10 peptides were mixed in the library with the equimolar concentration and the mix was added to each analyzed sample before in-gel trypsin digestion. Digested peptides were sequentially desalted using C18 Stagetips (3M Empore) and porous carbon material (TipTop Carbon, Glygen) as described in (Rappsilber et al., 2007) and resuspended in 15µl of 0.1% FA.

### Mass Spectrometry analysis with scheduled PRM method

To identify and measure the proportion of the histone modifications a parallel reaction monitoring method (RM) was used (Liebler and Zimmerman 2013). The mass spectrometer was operated in the scheduled RM mode to identify and quantify specific fragment ions of N-terminal peptides histone proteins. In this mode, the mass spectrometer automatically switched between one survey scan and 9 MS/MS acquisitions of the m/z values described in the inclusion list containing the precursor ions, modifications and fragmentation conditions. Survey full scan MS spectra (from m/z 270-730) were acquired with resolution 60,000 at m/z 400 (AGC target of 3×10^6). RM spectra were acquired with resolution 30,000 to a target value of 2 10^5^, maximum IT 60 ms, isolation window 0.7 m/z and fragmented at 27% or 30% normalized collision energy. Typical mass spectrometric conditions were: spray voltage, 1.5 kV; no sheath and auxiliary gas flow; heated capillary temperature, 250°C.

### Histone modifications quantification

Data analysis was performed with Skyline (version 3.7) (MacLean et al., 2010) by using doubly and triply charged peptide masses for extracted ion chromatograms (XICs). Selection of respective peaks was identified based on the retention time and fragmentation spectra of the spiked in heavy-labelled peptides. Integrated peak values (Total Area MS1) were exported as csv file for further calculations. Total area MS1 from endogenous peptides was normalized to the respective area of heavy-labelled peptides. The sum of all normalized total area MS1 values of the same isotopically modified peptide in one sample resembled the amount of total peptide. The proportions of the different K20 methylation states were calculated and displayed as percentages of the overall K20 peptide amount.

## Models

### HUA and mock models

We consider the proportions of un- (me0), mono- (me1), di- (me2) and tri-methylated (me3) H4K20 within a Xenopus embryo population, defined as

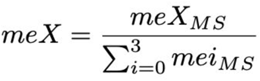

where meX_MS_ is the H4K20 methylation as measured by mass spectrometry and X ∈ {0,1,2,3}. We assume successive methylation and demethylation of H4K20 (van Nuland and Gozani, 2016) resulting in three possible methylation rate constants for mono-, di-, and tri-methylation with rate constants m_1_, m_2_, m_3_, respectively, and three possible demethylation rates with rate constants d_1_, d_2_, d_3_ (Figures 2A and 3A). However, reactions might share rate constants. The simplest model (Figure 2B left) comprises one shared methylation rate constant for mono-, di- and tri-methylation We successively added model-specific rate constants to this simplest model (Figure 2B). Models allowing for two specific methylation rate constants are identical to a model allowing for three specific methylation rate constants. Hence, we do not consider models with two specific methylation rate constants separately. This results in 2^3^-3 = 5 models for methylation - three methylation rate constants with either a shared or specific rate constant minus the three cases where we assume only two of the three rate constants to be specific. We have the same for demethylation resulting overall in (2^3^-3) *(2^3^-3) = 25 possible HUA models.

### Joint models

The joint model considers both mock and HUA data sets. We based the joint model on our previous findings assuming three specific methylation rate constants and at most one demethylation rate constant for both mock (Figure 2D) and HUA (Figure 3B) as well as a scaled Hill function with Hill coefficient 1 and offset 0.5 as cell-cycle function. In general, the joint model would allow for (2^3^-3)^4^ = 625 distinct models. By constraining both the HUA and mock model to allow for three methylation and one demethylation rate constants, we are able to reduce the number of possible models to 16. The simplest joint model is comprised of 3 rate constants which are shared for mock and the HUA reactions (Figure 4B left). We successively added model complexity by allowing for HUA-specific rate constants, totalling to 16 models for the joint model with demethylation in mock and HUA and 8 models for the joint model with demethylation present in either one or none (Figure 4B).

For all models we describe the temporal changes in these proportions by systems of ordinary differential equations (ODEs) using mass action kinetics (see below).

### HUA model

We first derive the system of ODEs for the absolute numbers of H4K20me states, given by 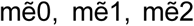, and 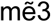, for the model with the largest number of rate constants (Figure 3A):

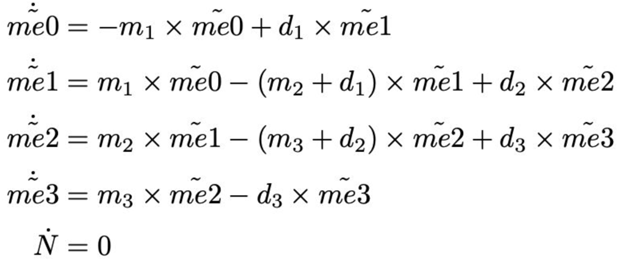

N is the total number of histone tails. As the HUA model assumes no cell-cycle, the number of histones over time is constant and its derivative is zero. The proportions me0, me1, me2 and me3 are given by 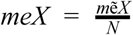, for X ∈ {0, 1, 2, 3} (Alabert et al., 2020) and the corresponding ODEs are given by

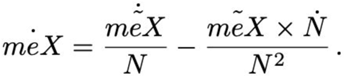

simplifying to

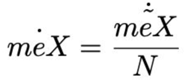

in the HUA model. The full ODE system for the proportions is given by

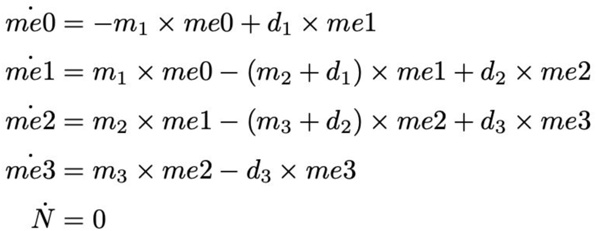

### Mock model - constant cell-cycle duration

According to the HUA model, we first formulate the ODE system of the absolute numbers of methylation states, 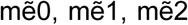, and 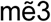. We assume for now the cell-cycle duration to be constant over time, denoted by a:

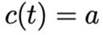

Then the full ODE system of the absolute numbers of methylation states is given by

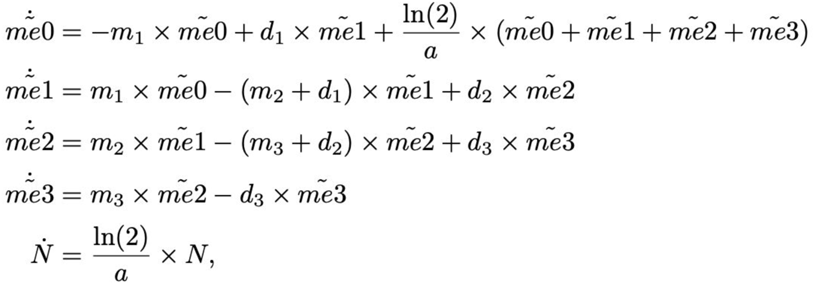

N is the total number of histone tails, where 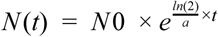 and *N* (*t*_0_) = *N*0 the number of histone tails at the beginning of the model. Then the ODE system of the methylation proportions is given by

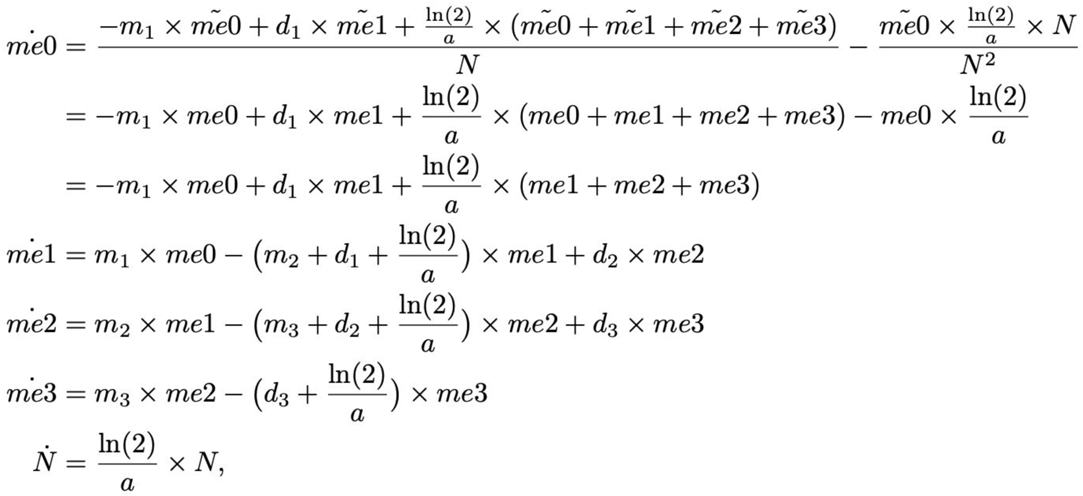

### Mock model - linearly increasing cell-cycle duration

Similarly, we derive the ODE system of the methylation proportions, me0, me1, me2 and me3, for linearly increasing cell-cycle function, where we first assumed the cell-cycle function to be

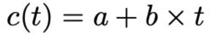

and hence,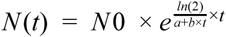

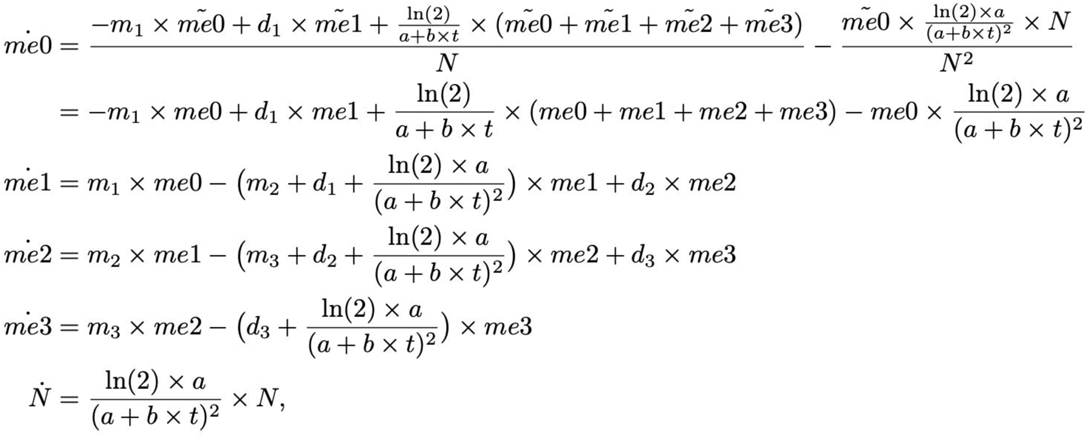

To constrain the system to biologically meaningful cell-cycle durations we included prior knowledge from literature: at 5.5 hpf the cell-cycle in Xenopus has been found to be ∼0.5 hours (Heasman, 2006). Hence, we assumed a second linearly increasing cell-cycle function

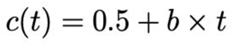

### Mock model - scaled Hill function with Hill coefficient 1 and offset as cell-cycle duration

Similarly, we derive the ODE system of the methylation proportions me0, me1, me2 and me3 for the cell-cycle function 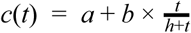 and 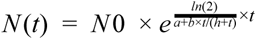:

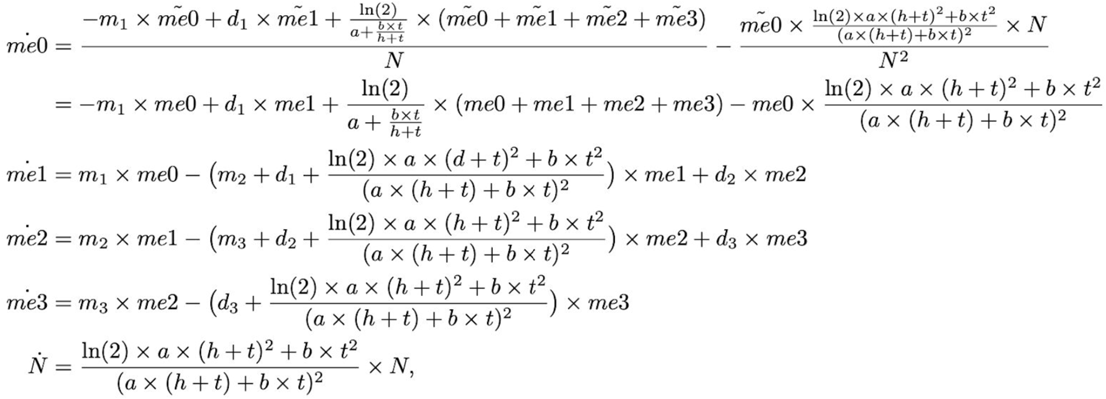

Similar to the mock model with linearly increasing cell-cycle function we tested three different scaled Hill functions with Hill coefficient 1 and offset as cell-cycle functions:

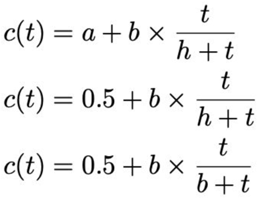

We again reduced the number of model parameters in the second equation by inserting prior knowledge about the cell-cycle duration at the start of the model (see Mock model - linearly increasing cell-cycle duration). Additionally, we reduced the number of model parameters further by assuming the scaling b and the dissociation constant h to be identical in the third equation. In comparison to the former two cell-cycle functions, the third equation constrains the parameter space more strictly. E.g. for upper and lower boundaries of 100 and 0.0001 for parameters a, b and h the first equation will allow for cell-cycle durations up to 100+100 *(42-5.5)/(0.0001+42-5.5) ≈ 200 hours while the third equation only allows for cell-cycle durations for up to 0.5+100 *(42-5.5)/(100+42-5.5) ≈ 27 hours.

### Noise models

As experimental data is generally noise corrupted, we evaluated all models with an underlying Laplacian noise model. Maier et. al. (Maier et al., 2017) have shown that Laplacian noise models may outperform Gaussian ones due to their increased robustness against outliers (Maier et al., 2017). All model parameters are comprised in the parameter vector *θ* and the experimental measurement *i* at time point *k* is denoted by 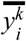. The log-likelihoods for the Laplacian noise model is given by

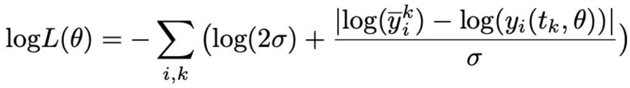

By performing maximum likelihood estimation we obtain the optimal model parameters.

### Optimization and parameter estimation

The model parameters include the relative initial states, me1_0_, me2_0_ and me3_0_, where me0_0_=0.1 is fixed to obtain structural identifiability and

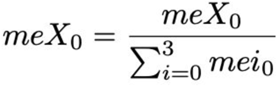

with X ∈ {0, 1, 2, 3}, one noise parameter, the model-specific rate constants of (de-)methylation and potentially up to three constants (mock models) describing the cell-cycle function. For numerical reasons we optimized the parameters in a log10 scale (Hass et al., 2019). The lower and upper bounds for the rate constants, initial states, noise parameter and cell-cycle parameters were initiated in log10 scale at −10 to 2, −4 to 2, −2 to 0 and −10 to 10, respectively. We performed multi-start local optimization of the negative log-likelihood using the parameter estimation toolbox ESTO (Stapor et al., 2018) and simulated the models with AMICI (Fröhlich et al., 2017). We performed at least 100 local optimization runs per model, initialized by latin-hyper cube-sampled starts. For the models not converging upon these initializations (where by ‘not converging’ we mean that the likelihood value of the second best run differs more than 0.1) we decreased the width between upper and lower bounds to increase the probability of convergence. For this, we assured that the optimization bounds were wide enough such that the optimal values are not in the bounds for the rate constants and the cell-cycle parameters. As the initial states are unidentifiable we ignored optimal values which ran into these boundaries as long as other optimal values were found within. For models where this was not the case we expanded the boundaries of the rate constants and initial states up to −20 to 10 and −10 to 10, respectively, as we assumed any smaller or larger values to be biologically non-informative. For the initially non-converged joint models we also increased the number of starts to 800. Models not having converged upon manually adjusting the boundaries and running for 800 starts were determined to not converge. All mock and HUA models converged. We determined 6 out of the 40 joint models to not converge (Supplementary Information). The given likelihood values of these joint models are lower bounds of the true optimal likelihood values obtainable upon convergence. As the likelihood values of all 6 non-converged joint models still resulted in considerably lower BIC values in comparison to the other tested models we can safely report them as best performing models for the respective demethylation hypothesis. As the BIC values between the demethylation hypotheses allowing and not allowing for demethylation in HUA differ considerably we assume the comparison between different demethylation hypotheses to be valid and the resulting conclusions to be justified.

### Model selection

We use the Bayesian Information Criterion (BIC) (Schwarz, 1978) for model comparisons,

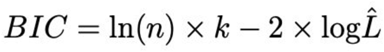

where *n* is the number of data points, *k* is the number of estimated parameters or the overall model complexity and log*L* is the log-likelihood value for the maximum likelihood estimate of the model parameters. The BIC rewards high likelihood values and penalizes model complexity. Hence, low BIC values are preferable. In comparison to other model selection methods such as the Akaike Information Criterion (AIC) the BIC penalizes additional model complexity more strongly. We consider a ΔBIC>10 between two models to be enough evidence to reject the model with the higher BIC (Kass and Raftery, 1995).

### Parameter uncertainty

To receive the uncertainties for the estimated model parameters we performed Markov Chain Monte Carlo (MCMC) sampling of the posterior distribution

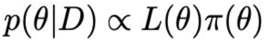

with uniform prior *π*(*θ*) defined over the optimization boundaries, likelihood function *L*(*θ*) and data *D*. We sampled the posterior for all six best performing joint models and the mock model with three specific methylation rate constants and no demethylation (ESTO-internal function *getParameterSamples*). We employed parallel tempering with five parallel chains initiated at the five most optimal parameter estimates per model obtained during optimization and performed 10^6^ iterations. Upon performing a Geweke test (first 10% versus last 50% of the final MCMC chains), we discarded the first 10% of the samples as burn-in phase and thinned the chains keeping only every 100^th^ sample. The marginal posterior distributions are plotted via violin plots (plotting function *violin*, Hoffman, H. (2015). violin.m - Simple violin plot using matlab default kernel density estimation. (https://de.mathworks.com/matlabcentral/fileexchange/45134-violin-plot), MATLAB Central File Exchange. Retrieved November 13, 2019.)).

### Validation - cell-cycle durations

We used the median and the 25^th^ and 75^th^ percentiles of the MCMC chain determined during the parameter uncertainty analysis for the cell-cycle parameter b, and evaluated the median and the 25^th^ and 75^th^ percentiles of the cell-cycle function according to

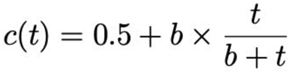

for *t* ∈ [0,40], where the cell-cycle duration of 0.5 hours at 5.5 hpf (start of model) is taken from (Anderson et al., 2017; Gelens et al., 2015).

### Prediction of number of cells

Using the median and the 25^th^ and 75^th^ percentiles of the cell-cycle parameter *b* (as determined in the validation analysis), we determined the theoretical number of cells a Xenopus embryo is on average composed of between 5.5 hpf and 45.5 hpf according to

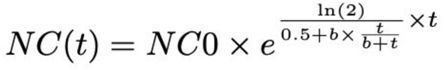

where NC(t) is the number of cells at time t, NC0 the initial number of cells and 0.5+b *t/(b+t) the cell-cycle function (constrained scaled Hill function with Hill coefficient 1 and offset 0.5). To account for the time-dependent cell-cycle function we calculated the cell numbers in time steps of 0.01 hours, where we take the initial cell number NC0(t) for every time step t to be the cell number of the previous time step NC(t-0.01). At the start of the model (at 5.5 hpf) we take the initial number of cells NC0(0) to be 4096 (Heasman, 2006).

### Implementation

The toolboxes used for the analysis of the manuscript for ODE simulation (AMICI (Fröhlich et al., 2017)) and parameter estimation (ESTO (Stapor et al., 2018)) are available under https://github.com/ICB-DCM. The analysis was performed with MATLAB 2017a.

## REFERENCES

Abbas, T., Shibata, E., Park, J., Jha, S., Karnani, N., and Dutta, A. (2010). CRL4(Cdt2) regulates cell proliferation and histone gene expression by targeting PR-Set7/Set8 for degradation. Mol. Cell 40, 9–21.

Alabert, C., Loos, C., Voelker-Albert, M., Graziano, S., Forné, I., Reveron-Gomez, N., Schuh, L., Hasenauer, J., Marr, C., Imhof, A., et al. (2020). Domain Model Explains Propagation Dynamics and Stability of Histone H3K27 and H3K36 Methylation Landscapes. Cell Rep. 30, 1223–1234.e8.

Anderson, G.A., Gelens, L., Baker, J.C., and Ferrell, J.E., Jr (2017). Desynchronizing Embryonic Cell Division Waves Reveals the Robustness of Xenopus laevis Development. Cell Rep. 21, 37–46.

Bannister, A.J., and Kouzarides, T. (2011). Regulation of chromatin by histone modifications. Cell Research 21, 381–395.

Barski, A., Cuddapah, S., Cui, K., Roh, T.-Y., Schones, D.E., Wang, Z., Wei, G., Chepelev, I., and Zhao, K. (2007). High-resolution profiling of histone methylations in the human genome. Cell 129, 823–837.

Centore, R.C., Havens, C.G., Manning, A.L., Li, J.-M., Flynn, R.L., Tse, A., Jin, J., Dyson, N.J., Walter, J.C., and Zou, L. (2010). CRL4Cdt2-Mediated Destruction of the Histone Methyltransferase Set8 Prevents Premature Chromatin Compaction in S hase. Molecular Cell 40, 22–33.

Feng, W., Yonezawa, M., Ye, J., Jenuwein, T., and Grummt, I. (2010). HF8 activates transcription of rRNA genes through H3K4me3 binding and H3K9me1/2 demethylation. Nat. Struct. Mol. Biol. 17, 445–450.

Fraga, M.F., Ballestar, E., Villar-Garea, A., Boix-Chornet, M., Espada, J., Schotta, G., Bonaldi, T., Haydon, C., Ropero, S., Petrie, K., et al. (2005). Loss of acetylation at Lys16 and trimethylation at Lys20 of histone H4 is a common hallmark of human cancer. Nat. Genet. 37, 391–400.

Fröhlich, F., Kaltenbacher, B., Theis, F.J., and Hasenauer, J. (2017). Scalable Parameter Estimation for Genome-Scale Biochemical Reaction Networks. PLoS Comput. Biol. 13, e1005331.

Gelens, L., Huang, K.C., and Ferrell, J.E., Jr (2015). How Does the Xenopus laevis Embryonic Cell Cycle Avoid Spatial Chaos? Cell Rep. 12, 892–900.

Graham, C.F., and Morgan, R.W. (1966). Changes in the cell cycle during early amphibian development. Developmental Biology 14, 439–460.

Greer, E.L., and Shi, Y. (2012). Histone methylation: a dynamic mark in health, disease and inheritance. Nat. Rev. Genet. 13, 343–357.

Harris, W.A., and Hartenstein, V. (1991). Neuronal determination without cell division in Xenopus embryos. Neuron 6, 499–515.

Hass, H., Loos, C., Raimúndez-Álvarez, E., Timmer, J., Hasenauer, J., and Kreutz, C. (2019). Benchmark problems for dynamic modeling of intracellular processes. Bioinformatics 35, 3073–3082.

Heasman, J. (2006). Patterning the early Xenopus embryo. Development 133, 1205–1217.

Jadhav, U., Manieri, E., Nalapareddy, K., Madha, S., Chakrabarti, S., Wucherpfennig, K., Barefoot, M., and Shivdasani, R.A. (2020). Replicational Dilution of H3K27me3 in Mammalian Cells and the Role of oised romoters. Mol. Cell.

Jambhekar, A., Dhall, A., and Shi, Y. (2020). Author Correction: Roles and regulation of histone methylation in animal development. Nat. Rev. Mol. Cell Biol. 21, 59.

Jasencakova, Z., Scharf, A.N.D., Ask, K., Corpet, A., Imhof, A., Almouzni, G., and Groth, A. (2010). Replication stress interferes with histone recycling and predeposition marking of new histones. Mol. Cell 37, 736–743.

Jørgensen, S., Schotta, G., and Sørensen, C.S. (2013). Histone H4 lysine 20 methylation: key player in epigenetic regulation of genomic integrity. Nucleic Acids Res. 41, 2797–2806.

Kass, R.E., and Raftery, A.E. (1995). Bayes Factors. Journal of the American Statistical Association 90, 773.

Klutstein, M., Nejman, D., Greenfield, R., and Cedar, H. (2016). DNA Methylation in Cancer and Aging. Cancer Research 76, 3446–3450.

Lachner, M., Sengupta, R., Schotta, G., and Jenuwein, T. (2004). Trilogies of Histone Lysine Methylation as Epigenetic Landmarks of the Eukaryotic Genome. Cold Spring Harbor Symposia on Quantitative Biology 69, 1–10.

MacLean, B., Tomazela, D.M., Shulman, N., Chambers, M., Finney, G.L., Frewen, B., Kern, R., Tabb, D.L., Liebler, D.C., and MacCoss, M.J. (2010). Skyline: an open source document editor for creating and analyzing targeted proteomics experiments. Bioinformatics 26, 966–968.

Maier, C., Loos, C., and Hasenauer, J. (2017). Robust parameter estimation for dynamical systems from outlier-corrupted data. Bioinformatics 33, 718–725.

Newport, J., and Kirschner, M. (1982). A major developmental transition in early xenopus embryos: I. characterization and timing of cellular changes at the midblastula stage. Cell 30, 675–686.

van Nuland, R., and Gozani, O. (2016). Histone H4 Lysine 20 (H4K20) Methylation, Expanding the Signaling Potential of the Proteome One Methyl Moiety at a Time. Mol. Cell. Proteomics 15, 755–764.

Oda, H., Okamoto, I., Murphy, N., Chu, J., Price, S.M., Shen, M.M., Torresadilla, M.E., Heard, E., and Reinberg, D. (2009). Monomethylation of Histone H4-ysine 20 Is Involved in Chromosome Structure and Stability and Is Essential for Mouse Development. Molecular and Cellular Biology 29, 2278–2295.

Pesavento, J.J., Yang, H., Kelleher, N.L., and Mizzen, C.A. (2008). Certain and progressive methylation of histone H4 at lysine 20 during the cell cycle. Mol. Cell. Biol. 28, 468–486.

Rappsilber, J., Mann, M., and Ishihama, Y. (2007). rotocol for micro-purification, enrichment, pre-fractionation and storage of peptides for proteomics using StageTips. Nat. Protoc. 2, 1896–1906.

Reverón-Gómez, N., González-Aguilera, C., Stewart-Morgan, K.R., Petryk, N., Flury, V., Graziano, S., Johansen, J.V., Jakobsen, J.S., Alabert, C., and Groth, A. (2018). Accurate Recycling of Parental Histones Reproduces the Histone Modification Landscape during DNA Replication. . Cell 72, 239–249.e5.

Sabherwal, N., Thuret, R., Lea, R., Stanley, P., and Papalopulu, N. (2014). aPKC phosphorylates p27Xic1, providing a mechanistic link between apicobasal polarity and cell-cycle control. Dev. Cell 31, 559–571.

Sakaguchi, A., and Steward, R. (2007). Aberrant monomethylation of histone H4 lysine 20 activates the DNA damage checkpoint in Drosophila melanogaster. J. Cell Biol. 176, 155–162.

Sanders, S.L., Portoso, M., Mata, J., Bähler, J., Allshire, R.C., and Kouzarides, T. (2004). Methylation of Histone H4 Lysine 20 Controls Recruitment of Crb2 to Sites of DNA Damage. Cell 119, 603–614.

Schneider, T.D., Arteaga-Salas, J.M., Mentele, E., David, R., Nicetto, D., Imhof, A., and Rupp, R.A.W. (2011). Stage-specific histone modification profiles reveal global transitions in the Xenopus embryonic epigenome. PLoS One 6, e22548.

Schotta, G., Lachner, M., Sarma, K., Ebert, A., Sengupta, R., Reuter, G., Reinberg, D., and Jenuwein, T. (2004). A silencing pathway to induce H3-K9 and H4-K20 trimethylation at constitutive heterochromatin. Genes Dev. 18, 1251–1262.

Schotta, G., Sengupta, R., Kubicek, S., Malin, S., Kauer, M., Callén, E., Celeste, A., Pagani, M., Opravil, S., De La Rosa-Velazquez, I.A., et al. (2008). A chromatin-wide transition to H4K20 monomethylation impairs genome integrity and programmed DNA rearrangements in the mouse. Genes Dev. 22, 2048–2061.

Schwarz, G. (1978). Estimating the Dimension of a Model. The Annals of Statistics 6, 461–464.

Southall, S.M., Cronin, N.B., and Wilson, J.R. (2013). A novel route to product specificity in the Suv4-20 family of histone H4K20 methyltransferases.

Stapor, P., Weindl, D., Ballnus, B., Hugß, S., Loos, C., Fiedler, A., Krause, S., Hrof1, S., Fröhlich, F., Hasenauer, J., et al. (2018). PESTO: arameter EStimation TOolbox. Bioinformatics 34, 705–707.

Thuret, R., Auger, H., and Papalopulu, N. (2015). Analysis of neural progenitors from embryogenesis to juvenile adult in Xenopus laevis reveals biphasic neurogenesis and continuous lengthening of the cell cycle. Biol. Open 4, 1772–1781.

Villar-Garea, A., Forne, I., Vetter, I., Kremmer, E., Thomae, A., and Imhof, A. (2012). Developmental regulation of N-terminal H2B methylation in Drosophila melanogaster. Nucleic Acids Res. 40, 1536–1549.

Xiao, B. (2005). Specificity and mechanism of the histone methyltransferase Pr-Set7. Genes & Development 19, 1444–1454.

Yang, H., Pesavento, J.J., Starnes, T.W., Cryderman, D.E., Wallrath, L.L., Kelleher, N.L., and Mizzen, C.A. (2008). Preferential dimethylation of histone H4 lysine 20 by Suv4-20. J. Biol. Chem. 283, 12085–12092.

Zee, B.M., Britton, L.-M.L., Wolle, D., Haberman, D.M., and Garcia, B.A. (2012). Origins and formation of histone methylation across the human cell cycle. Mol. Cell. Biol. 32, 2503–2514.

